# A phased chromosome-level genome of the annelid tubeworm *Galeolaria caespitosa*

**DOI:** 10.1101/2024.11.18.624032

**Authors:** Monique van Dorssen, Emily K. Belcher, Cristóbal Gallegos, Kathryn A. Hodgins, Keyne Monro

## Abstract

Haplotype-resolved (phased) genome assemblies are emerging as important assets for genomic studies of species with high heterozygosity, but remain lacking for key animal lineages. Here, we use PacBio HiFi and Omni-C technologies to assemble the first phased, annotated, chromosome-level genome for any annelid: the reef-building tubeworm *Galeolaria caespitosa* (Serpulidae). The assembly is 803.5 Mbp long (scaffold N50 = 76.5 Mbp) for haplotype 1 and 789.3 Mbp long (scaffold N50 = 75.4 Mbp) for haplotype 2, which are arranged into 11 pairs of chromosomes showing no sign of sex chromosomes. This compares with cytological analyses reporting 12–13 pairs in *Galeolaria*’s closest relatives, including species that are protandrous hermaphrodites. We combined long-read and short-read transcriptome sequencing to annotate both haplotypes, resulting in 43,191 predicted proteins for haplotype 1, 39,675 proteins for haplotype two, and 55.5% of proteins with at least one functional annotation. We also assembled a mitochondrial genome 23 Kbp long, annotating all genes typically found in mitochondrial DNA apart from those coding the *16S* ribosomal subunit and the protein *atp8* — a short, fast-evolving mitochondrial gene missing in other metazoans. Comparing *Galeolaria*’s genome to those of three other annelids reveals limited collinearity despite 32.2% of shared orthologous gene clusters (4,248 of 13,174 clusters counted in *Galeolaria*), suggesting extensive chromosomal rearrangements among lineages. New high-quality annelid genomes may help resolve the genetic and evolutionary basis of this diversity.

## Introduction

Genomic tools are increasingly used to understand biodiversity at the genetic level, but remain out of reach for most non-model species of eco-evolutionary or conservation concern due to a lack of high-quality reference genomes (Formenti et al., 2022; Paez et al., 2022). Such genomes can greatly enhance genomic insights by providing templates on which to map genomic information and study its broader function (Hoban et al., 2016; Mérot et al., 2020). To date, however, most genome assemblies are either in draft form (i.e., fragmented and unannotated), or merge the two parental haplotypes present in the sequenced individual (Duitama, 2023). Although chromosome-level assemblies are fast accumulating, haplotype-merged ones may only be accurate for inbred model species, or species with low heterozygosity, where haplotypes are genetically similar (Takeuchi et al., 2022). For outbred, highly heterozygous species, however, merging haplotypes may hide substantial variation owing to differences between them, and distort our view of genetic diversity.

Marine invertebrates, and broadcast spawners especially, are known for high levels of heterozygosity driven by large population sizes, high fecundities, and extensive dispersal of gametes, embryos, and larvae (Lotterhos et al., 2021; Plough, 2016). Moreover, by representing the majority of extant animal phyla, including the oldest metazoan lineages, marine invertebrate genomes may encode not only unique gene products, but also key insights about adaptation and diversification (Lopez et al., 2019). Genome assemblies that are phased into haplotypes, and capture information about heterozygosity, are therefore essential to unlock and study genetic diversity in this group. Despite the challenges that high levels of heterozygosity bring to the assembly of such genomes, as the presence of divergent alleles can lead to fragmented assemblies or misassembles, advances in long-read sequencing technologies now enable these in the absence of parental sequencing or other pedigree information (Cheng et al., 2021; Cheng et al., 2022). Currently though, only a handful of phased or well-annotated genomes exist for marine invertebrates, covering only a small sample of phyla such as molluscs (Takeuchi et al., 2022) and echinoderms (Schiebelhut et al., 2023).

Annelids remain one of the most diverse, yet poorly understood, metazoan phyla (Sun et al., 2021). They have evolved a remarkable array of reproductive and developmental modes (Dorresteijn & Westheide, 1999), and play major roles in the functioning of benthic communities from shallow waters to deep-sea trenches (Hutchings, 1998). Annelid genomes could thus provide important new insights into life history evolution and adaptation to environmental change. Despite the biodiversity captured in this group, we know surprisingly little about the genomic basis of annelid life histories or their evolutionary diversification (Lopez et al., 2019; Sun et al., 2021). Of the 36 chromosome-level genome assemblies currently available for annelids in NCBI (accessed June 2024), few are published (but see Jin et al., 2020; Zakas et al., 2022) and none appear to be both phased and annotated. Such oversight leaves major gaps in our understanding of basic evolutionary biology and genome evolution in a key metazoan lineage, and limited tools for progress on this front.

Here, we present the first phased and annotated chromosome-level genome for any annelid — the reef-building tubeworm, *Galeolaria caespitosa. G. caespitosa* is a broadcast-spawning species endemic to rocky shores of temperate Australia, where it is a coastal ecosystem engineer (Cole et al., 2018; Wright & Gribben, 2017) and bioindicator of ecosystem health (Lu et al., 2017; Ross & Bidwell, 2001). *Galeolaria* also belongs to the Serpulidae, an ecologically-important family that includes notorious biofoulers and bioinvaders (Sun et al., 2021), and diverse life histories spanning different modes of sex determination, reproduction, and development (Kupriyanova et al., 2001). Little is known about the genomic basis of these patterns, despite several lines of evidence linking them to changes in chromosome number (Dasgupta & Austin, 1960), structural rearrangements within chromosomes (Dixon et al., 1998), and accelerated substitution rates and structural rearrangements within mitochondrial genomes (Sun et al., 2021). As the first annelid genome of this quality, the *Galeolaria* genome will be valuable in its own right, and serve as a reference for comparison with other individuals, species, and metazoan lineages.

## Methods

### DNA extraction, library preparation, and sequencing

We assembled the genome of a wild male, sampled from Chelsea Pier (Victoria, Australia) at low tide. The male was removed from its tube to induce spawning, then sperm and muscle tissue were frozen separately in liquid nitrogen and stored at -80 °C until DNA extraction.

High-molecular-weight genomic DNA was extracted from sperm using Qiagen’s Blood & Cell Culture DNA Kit. A PacBio HiFi library was then prepared using PacBio’s SMRTbell Express Template Prep Kit, and sequenced in two SMRT Cells on the PacBio Sequel II platform (Cantata Bio, California, USA).

A Dovetail Omni-C library was prepared from muscle tissue, using Dovetail’s Omni-C Kit. Chromatin was fixed in place in the nucleus using formaldehyde, before digestion with DNAse I. Chromatin ends were repaired and ligated to a biotinylated bridge adapter, followed by proximity ligation of adapter-containing ends and reversal of crosslinks. DNA was then purified and treated to remove biotin outside ligated fragments. The library was constructed using NEBNext Ultra enzymes and an Illumina-compatible adapter, with biotin-containing fragments isolated by streptavidin beads before PCR enrichment, before sequencing on the Illumina HiSeq X platform (Cantata Bio, California, USA).

### Initial scaffold-level assembly

We estimated *k*-mer counts (*k* = 21) for Omni-C reads using *jellyfish* v2.3.0 (Marçais & Kingsford, 2011), *k*-mer completeness using *Merqury* v1.3 (Rhie et al., 2020), and genome size and heterozygosity using *GenomeScope* v2.0 (Ranallo-Benavidez et al., 2020) with default settings (Supplementary Figure 1B).

**Figure 1.**
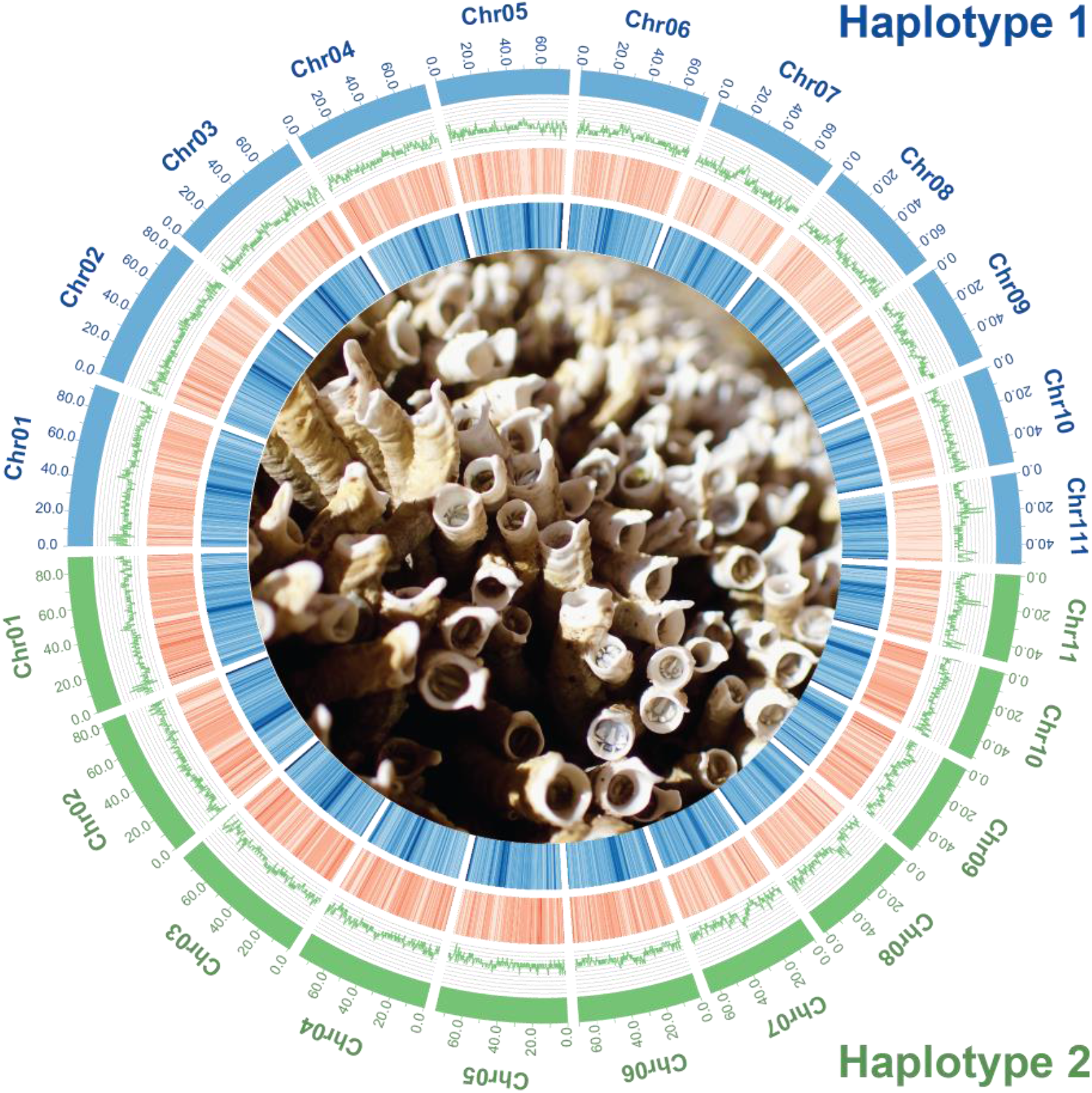
Circos plot of the 11 chromosomes in the haplotype-resolved assembly. From outside to inside: chromosomes of haplotype 1 (blue) and haplotype 2 (green), GC%, gene density (red), repeat density (blue), and *Galeolaria caespitosa* adults retracted into tubes at low tide (photo credit: E. Chirgwin). GC%, gene density, and repeat density were estimated in 500 Kb windows with a 100 Kb step.

To obtain phased contigs (haplotype 1 and haplotype 2) from the HiFi reads, we used *Hifiasm* v0.15.4-r347 (Cheng et al., 2021) in “*Hi-C Intergrated-Assembly*”-mode using the Omni-C reads for the phasing. We then scaffolded the phased contigs with the Omni-C reads using *YaHS* v1.1 (Zhou et al., 2022), calculated genome summary statistics for each scaffold-level assembly using *QUAST* v.5 (Mikheenko et al., 2018), and checked contiguity visually using chromatin contact-maps (Supplementary Figure 1A) made in *Juicebox* v1.9.8 (Durand et al., 2016).

We checked the completeness of assemblies using *BUSCO* v5.1.3 (Simão et al., 2015) queried against the eukaryote database (*eukaryota_odb10*, created January 2024 with 70 genomes and 255 BUSCOs).

### Chromosome-level genome assembly

We identified chromosomes from chromatin contact maps, which assemble chromosome-level scaffolds by translating the proximity of genomic regions in 3D-space to contiguous linear organisation. To test if the number of chromosomes identified from maps corresponded to a drop in scaffold size, we performed a clustering analysis on the lengths of the 20 longest scaffolds using the *k-means++* method (Arthur & Vassilvitskii, 2007) implemented using *SciStatCalc* (https://scistatcalc.blogspot.com/). After selecting the 11 largest scaffolds as chromosomes, we aligned the two haplotypes using *minimap2* v2.16 (Li, 2018), plotted the alignments in R using *ggpubr* (Kassambara, 2022) and *pafr* (Winter et al., 2022), and created dot-plots from *paf*-files containing the alignments (Supplementary Figure 1C). This allowed us to identify chromosomes that matched between haplotypes, and rename chromosomes accordingly. We then moved the largest version of each chromosome into haplotype 1 and the other version into haplotype 2, meaning these haplotypes may differ from those at the scaffold level above.

We re-calculated genome summary statistics for chromosome-level assemblies with *QUAST, Merqury*, and *BUSCO*, as above.

### Mitochondrial genome assembly

We extracted an initial mitochondrial assembly from the scaffold-level haplotype 1 using the *MitoHifi* pipeline (https://github.com/marcelauliano/MitoHiFi; Uliano-Silva et al., 2023), with *MITOS* (Bernt et al., 2013) as the annotation tool and the mitochondrial genome of the serpulid, *Hydroides elegans* (Zeng & Wang, 2022), as the starting reference. Since several expected genes were missing, a second annotation was performed with *MitoFinder* (Allio et al., 2020). The mitochondrial genome of *G. caespitosa* was plotted alongside that of *H. elegans* and *Spirobranchus giganteus* (Seixas et al., 2017) using circularMT (Goodman & Carr, 2024, Figure 2A).

**Figure 2.**
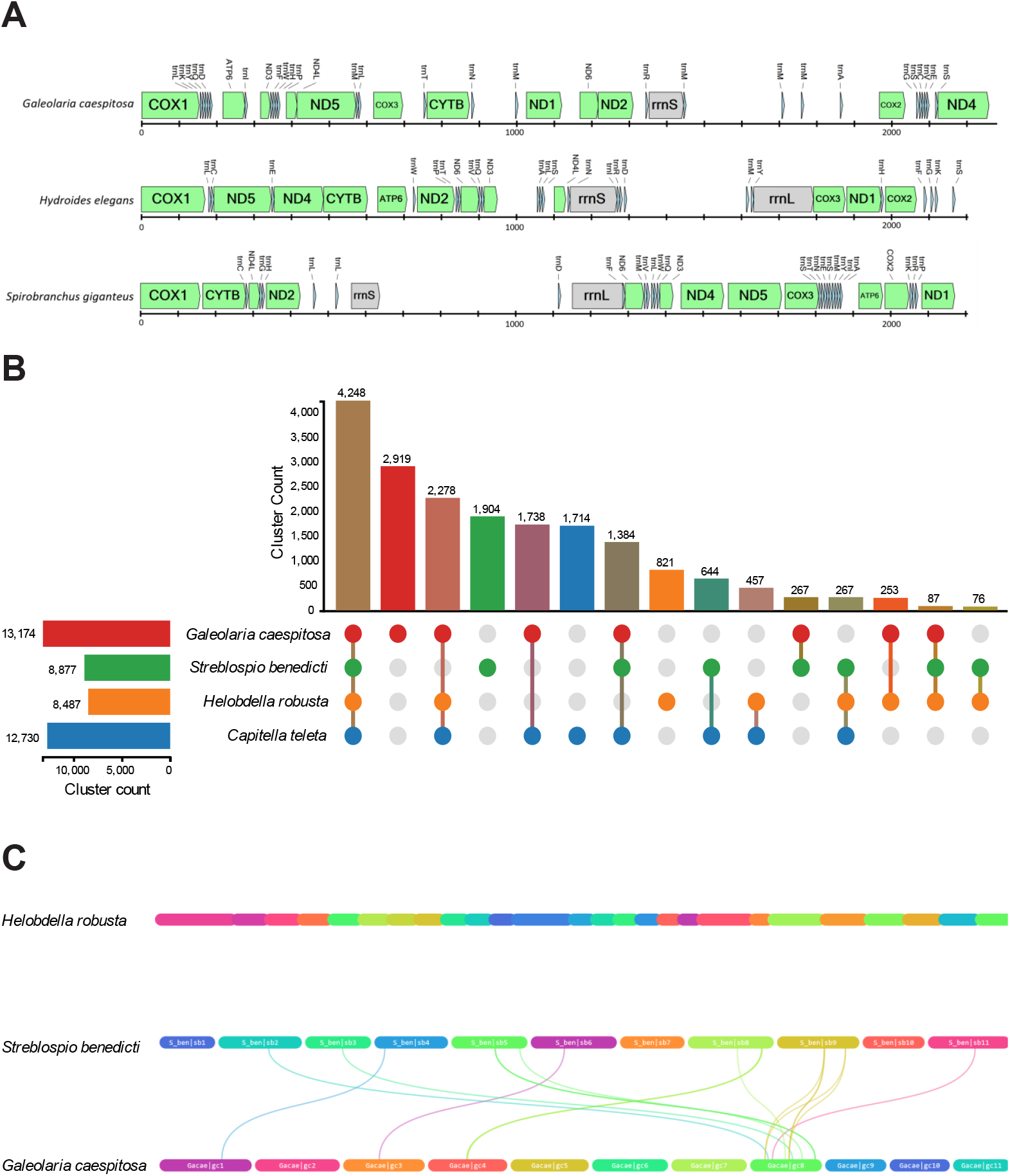
Comparative analyses of annelid genomes. (A) Mitochondrial gene order of *Galeolaria caespitosa, Hydroides elegans*, and *Spirobranchus giganteus*. (B) Orthologous cluster counts between *Galeolaria caespitosa, Streblospio benedicti, Helobdella robusta*, and *Capitella teleta*. (C) Collinearity analysis comparing *Helobdella robusta, Streblospio benedicti*, and *Galeolaria caespitosa*. Note that no synteny was detected between the genome of *Helobdella* and those of other species, and genomes of other species are labelled by chromosome number.

### Transcriptome sequencing for annotation

Both long-read and short-read transcriptomes were used for genome annotation. We generated full-length transcripts from pooled adults (also from Chelsea Pier) and developmental stages obtained by spawning adults and cross-fertilising gametes (Gallegos et al., 2024; Rebolledo et al., 2023). Developing embryos and larvae were sampled at appropriate times (~24 hours to 12 days) and stored at -80 °C until RNA extraction. We generated short-read transcripts from 91 samples of embryos obtained by spawning and cross-fertilising additional sets of pooled adults (from Brighton Pier, Victoria, Australia). Total RNA was extracted separately from adult males, adult females, unfertilized eggs, embryos, early larvae, and advanced larvae. Extraction required optimization by life stage, but used Trizol, a PureLink RNA Mini Kit with or without added Trizol, or a Qiagen RNeasy Plus Mini Kit with added mercaptoethanol, following manufacturer instructions.

RNA for long-read transcripts was pooled across life stages, underwent PacBio Iso-Seq library preparation using PacBio’s SMRTbell Express Template Prep Kit, and was sequenced on the PacBio Sequel IIe platform (Azenta Life Sciences, GENEWIZ Genomics Centre, Suzhou, China). Primers were removed and poly(A) tails trimmed from transcripts using *lima* v2.9.0 and *isoseq3* v4.0.0, respectively, from the PacBio Iso-Seq pipeline (PacBio, 2018). RNA for short-read transcripts underwent Illumina Stranded mRNA library preparation with poly(A) enrichment, and was sequenced on the NovaSeq X Plus platform (Australian Genome Research Facility, Melbourne, Australia). Adapters and low-quality sequences were removed transcripts using *fastp* v0.20.0 (Chen et al. 2018; Chen 2023).

### Structural annotation and gene models

Structural annotation was performed on both chromosome-level haplotypes. To improve gene-model predictions, we softmasked repeats in both assemblies before annotation using *RepeatModeler2* (Flynn et al., 2020) and *RepeatMasker* v4.1.1(Smit et al., 2013-2015). All known annelid proteins (151,839 in the UniProtKB database; The UniProt Consortium, 2022) were excluded from repeats using *ProtExcluder* v1.2 (Campbell et al., 2013). Repeat modeling also allowed us to characterize repeat type and density for each haplotype.

We predicted gene models and generated a structural annotation for each haplotype by combining Iso-Seq and RNA-seq evidence with evidence from annelid proteins downloaded from UniProtKB. First, we mapped Iso-Seq reads to each haplotype using *minimap2* v2.17 (Li, 2018), sorted them using *samtools* v1.9 (Danecek et al., 2021), collapsed them using *isoseq3* v4.0.0 (PacBio, 2018), and assembled the transcriptome using *stringtie* v2.1.5 (Pertea et al., 2015). RNA-seq reads were mapped to haplotypes using *STAR* v2.7.11 (Dobin et al., 2013). Next, following the BRAKER long-read integration protocol (Hoff et al., 2019; Stanke et al., 2008; Stanke et al., 2006), we annotated genes using BRAKER1 (Hoff et al., 2016) on mapped RNA-seq reads, BRAKER2 (Brůna et al., 2021) on UniProtKB protein sequences, and GeneMarkS-T (Tang et al., 2015) on mapped Iso-seq reads. Last, we used TSEBRA (Gabriel et al., 2021) to combine the three gene sets based on all extrinsic evidence. We used the long-read configuration and increased the weights of short-read and long-read hint sources, as recommended when the protein database lacks close relatives. We also reduced the threshold for intron support to 0.2, as recommended by TSEBRA. This produced a *gtf* file with structural annotations and gene models.

We converted the *gtf* file to a *gff3* file using AGAT v1.0.0 (Dainat, 2023), then cleaned the *gff3* file using *GFFtk* v24.2.4 (https://github.com/nextgenusfs/gfftk) and extracted protein sequences using *gffread* v0.12.7. To check the completeness our final protein set, we used *BUSCO* v5.1.3 queried against the eukaryote (*eukaryota_odb10*) database as above.

### Functional annotation

Functional annotation was also performed on both chromosome-level haplotypes by adapting the pipeline of Santangelo et al. (2023; https://github.com/James-S-Santangelo). We added functional annotations by querying predicted proteins against various databases (listed below), before merging and formatting annotations for upload to NCBI. First, we retrieved functional annotations using *InterProScan* v5.61.93-0 (Jones et al., 2014) and *Eggnog-mapper* v2.1.10 (Cantalapiedra et al., 2021). The resulting outputs were passed as input to *funannotate* v1.8.14 (Palmer & Stajich, 2019), which combined the annotations and queried additional databases (*merops* v12.0, *uniprot* v2024-01, *dbCan* v12.0, *pfam* v 36.0, *repeats* v.1.0, *go* 2024-01-17, *mibig* v1.4, *interpro* v98.0, *busco_outgroups* v1.0 metazoa database, and *gene2product* v.1.92). For any protein annotated as hypothetical and containing a fully-resolved 4-digit Enzyme Commission (EC) number, we replaced the annotation with the EC number’s product in the *ExPASSY* Enzyme database (Bairoch, 2000). Finally, we blasted protein-coding gene models against the NCBI *nr* database using *Diamond-blastp* v1.19 (Buchfink et al., 2021), and added results to the final *gff3* file using *gffutils* v0.13. We checked the completeness our final annotated protein set using *BUSCO* v5.1.3 queried against the eukaryote (*eukaryota_odb10*) database as above.

### Comparison with other annelid genomes

We compared the haplotype 1 protein set to three other annelid genomes using the orthologous and collinearity analyses of OrthoVenn3 (Sun et al., 2023), implemented with default parameters in the online module. The other genomes were the marine annelids, *Streblospio benedicti* (the closest relative with a comparable published assembly; Zakas et al., 2022) and *Capitella teleta*, and the leech *Helobdella robusta* (with the latter two genomes available in the OrthoVenn3 database). The collinearity analysis omitted the *Capitella* genome since no annotation was available. Last, to compare the 11 chromosomes identified in our assembly and the *Streblospio* assembly, we performed a reciprocal blast between their protein sequence files using *BLAST+* v12.15.0 (Camacho et al., 2009) to generate input for a pairwise synteny search using *MCScanX* (Wang et al., 2012), which we ran both with the default settings, and reducing the number of genes required to call synteny to 3 (default 5). We further generated completeness metrics for each of these genomes (including protein sets) using *BUSCO* v5.1.3 (Simão et al., 2015).

## Results

### Raw sequencing output

PacBio HiFi sequencing produced 58.6 Gbp of reads. Summary statistics from *GenomeScope* (Supplementary Figure 1B) estimated a genome ~707 Mb long, containing 43.7% of repetitive sequences and 4.8% heterozygosity.

### Initial scaffold-level genome assembly

The initial scaffold-level assembly had a total length of 904 Mbp and N50 of 24 Mb. Chromatin contact maps of scaffold-level haplotypes indicated that both genomes were highly contiguous (Supplementary Figure 1A). Haplotype 1 had 583 scaffolds with a total length of 803.5 Mb and *BUSCO* completeness score of 95.3%. Haplotype 2 had 289 scaffolds with a total length of 789.3 Mb and *BUSCO* completeness score of 96.1%. Total *k*-mer completeness was 99.4%. See Table 2 for full assembly statistics.

**Table 1.**
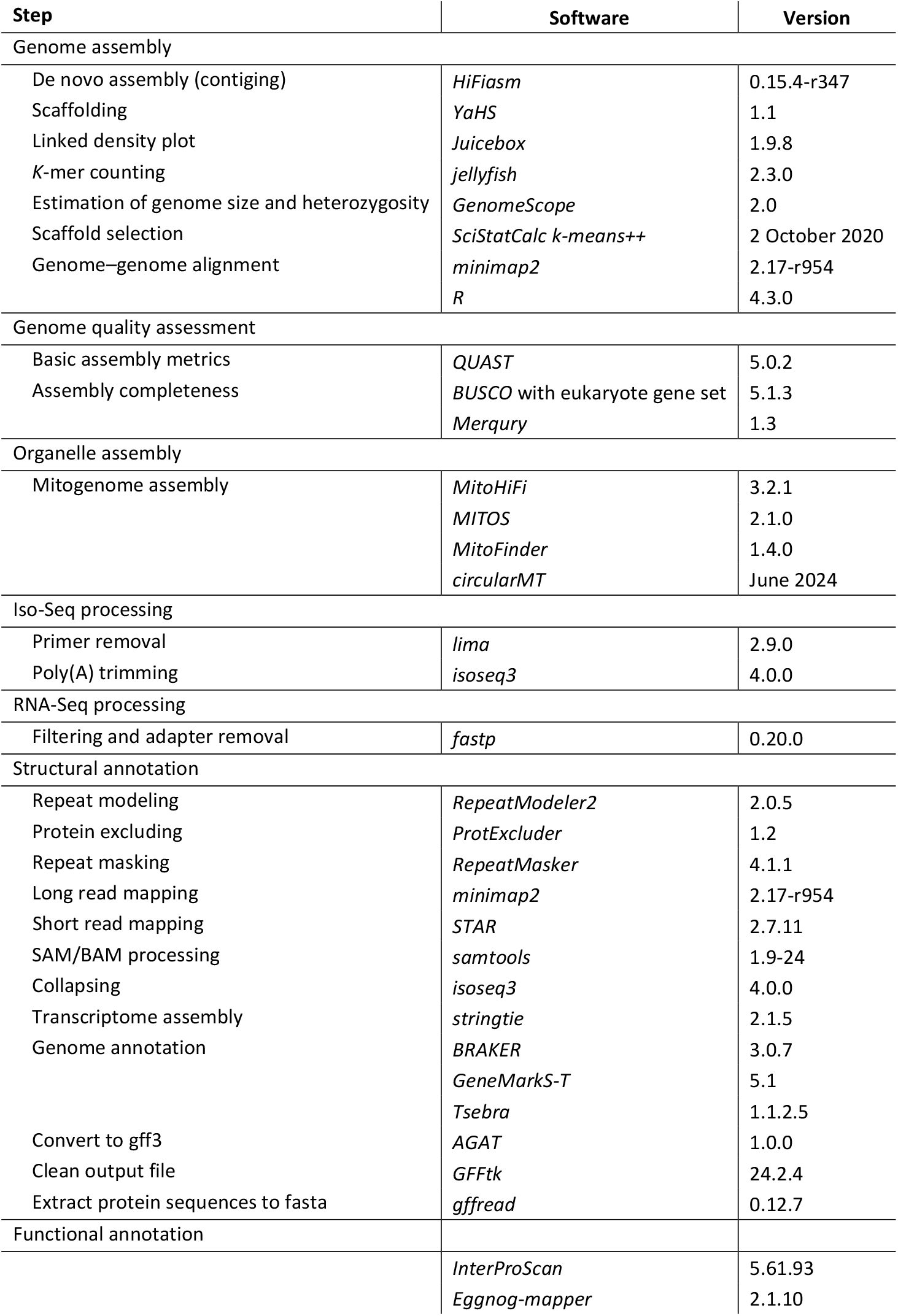
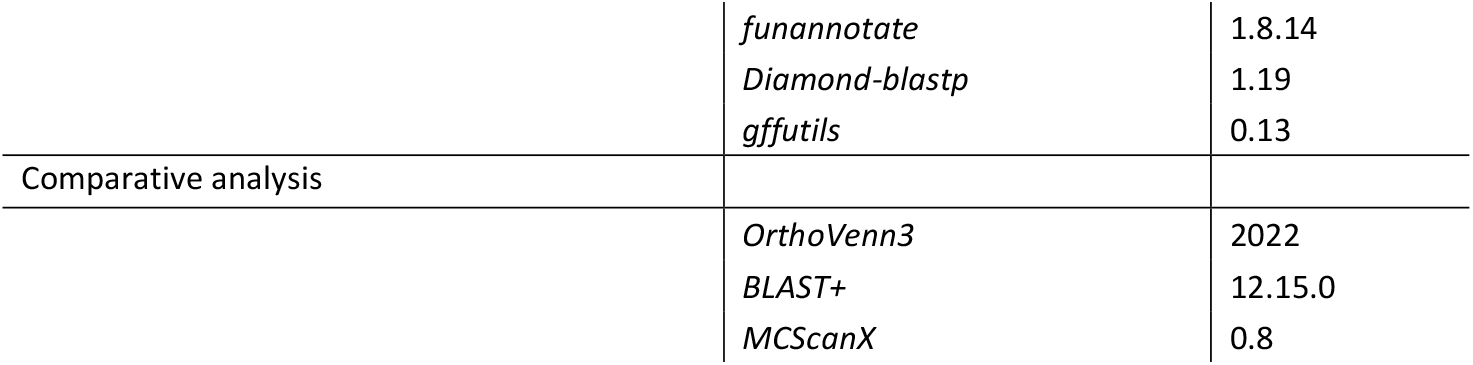
Assembly software and pipeline. Citations are listed in the text and scripts are available at https://github.com/moniquevdor/GaleolariaReferenceGenome.

**Table 2.** Sequencing and assembly statistics calculated with *QUAST* v5 and *BUSCO* v5.1.3 querying against the eukaryote database (*eukaryota_odb10*, created January 2024 with 70 genomes and 255 BUSCOs).

|  | Haplotype 1 assembly<br>(scaffold-level) | Haplotype 1 assembly<br>(chromosome-level) | Haplotype 2 assembly<br>(scaffold-level) | Haplotype 2 assembly<br>(chromosome-level) |
| --- | --- | --- | --- | --- |
| Number of scaffolds | 583 | 11 | 289 | 11 |
| Number of contigs | 676 | - | 381 | - |
| Total length (bp) | 803,490,011 | 792,440,614 | 789,320,953 | 759,533,915 |
| Largest scaffold (bp) | 89,960,646 | 92,870,682 | 92,870,682 | 89,960,646 |
| Largest contig (bp) | 33,040,180 | - | 33,497,296 | - |
| Number of Ns per 100 kbp | 2.61 | 2.52 | 2.84 | 2.9 |
| Scaffold N50 (bp) | 76,458,570 | 76,458,570 | 75,412,870 | 75,412,870 |
| Scaffold L50 | 5 | 5 | 5 | 5 |
| Contig N50 (bp) | 11,706,541 | - | 11,345,269 | - |
| Contig L50 | 22 | - | 22 | - |
| auN | 70,536,478 | 74,107,031 | 71,257,419 | 71,342,592 |
| Scaffold N90 (bp) | 55,310,200 | 55,531,921 | 52,145,674 | 52,145,674 |
| Scaffold L90 | 10 | 10 | 10 | 10 |
| Contig N90 (bp) | 2,789,309 | - | 3,633,209 | - |
| Contig L90 | 73 | - | 72 | - |
| GC (%) | 34.53% | 34.39% | 34.48% | 34.4% |
| Assembly BUSCO scores |  |  |  |  |
| Core genes queried | 255 | 255 | 255 | 255 |
| Complete BUSCOs (C) | 95.3% | 96.1% | 96.1% | 95.3% |
| Single-copy BUSCOs (S) | 94.3% | 95.7% | 94.8% | 95.3% |
| Duplicated BUSCOs (D) | 0.4% | 0.4% | 1.2% | 0.0% |
| Fragmented BUSCOs (F) | 4.3% | 3.5% | 3.5% | 4.3% |
| Missing BUSCOs (M) | 0.4% | 0.4% | 0.4% | 0.4% |
| Annotation stats |  |  |  |  |
| Number of predicted proteins | - | 43,191 | - | 39,675 |
| Number of genes with at least one functional annotation | - | 23,572 | - | 22,459 |
| Annotation BUSCO scores |  |  |  |  |
| Core genes queried | - | 255 | - | 255 |
| Complete BUSCOs (C) | - | 93.0% | - | 94.5% |
| Single-copy BUSCOs (S) | - | 92.2% | - | 94.1% |
| Duplicated BUSCOs (D) | - | 0.8% | - | 0.4% |
| Fragmented BUSCOs (F) | - | 5.9% | - | 3.9% |
| Missing BUSCOs (M) | - | 1.1% | - | 1.6% |

### Chromosome-level genome assembly

Chromatin contact maps showed scaffolds to be assembled into 11 major bins, supporting arrangement of the nuclear genome into 11 major chromosomes (Supplementary Figure 1A). *K*-means clustering of scaffold lengths reiterated this arrangement by clustering scaffolds 1-11 together (Supplementary Figure S2). Both the contact maps and dot-plot alignment of haplotypes (Supplementary Figures 1A and 1C) identified three potential inversions: a large inversion on chromosome 2 and two smaller inversions on chromosomes 9 and 11. At this level, haplotype 1 was 792.4 Mb long with a *BUSCO* completeness score of 96.1%, and haplotype 2 was 759.3 Mb long with a *BUSCO* completeness score of 95.3%. Total *k*-mer completeness was 99.3%. Full assembly statistics are again reported in Table 2.

We softmasked ~48% of each haplotype (~379 Mbp of haplotype 1 and ~362 Mbp of haplotype 2), then characterized repeat type and density for both of them (Figure 1, Supplementary Table S1). A large proportion (~36.5%) of repetitive sequences were unclassified, consistent with Zakas et al. (2022). Of the classified repetitive sequences, most (8.1% of haplotype 1 and 8.6% of haplotype 2) were long terminal repeats.

### Mitochondrial genome assembly

Using *MITOS*, the mitochondrial genome was assembled into a circular contig 22,809 bp long and containing 38 genes (Supplementary Table S2). Of these genes, 11 coded proteins, 26 coded transfer RNAs (tRNAs), and 1 coded *12S* ribosomal RNA (rRNA). We retrieved all common protein-coding genes except *nd3* and *atp8*, and all common tRNA genes (including single and double tRNA genes and 5 copies of the *tRNA-Met* gene), but were unable to identify the *16S* rRNA gene.

Using *MitoFinder*, we were able to identify *nd3*, but not 6 other protein-coding genes (*cox2, nd2, nd4, nd6*, and *atp6*, and *atp8*). We were also able to identify 5 copies of the *tRNA-Met* gene, 4 copies of the *tRNA-Glu* gene, and 2 copies of the *tRNA-Gly* gene, but neither of the rRNA genes. We therefore completed the mitochondrial genome by extracting the gene sequence for *nd3* from the *MitoFinder* genbank file, and performing a *blastn* search to identify its position in the *MITOS fasta* file. We then added *nd3* with its correct position to the *MITOS gff* file, producing a final circular genome with 39 fully-annotated genes. The alignment between the mitochondrial genomes of *Galeolaria caespitosa, Hydroides elegans*, and *Spirobranchus giganteus* revealed major gene rearrangements across species (Figure 2A), and *atp8* appeared to be missing from all three of these genomes.

### Transcriptome sequencing and genome annotation

Long-read transcriptome sequencing yielded 508,317 transcripts with a total length of 917 Mbp (N50 = 2,139 bp), maximum length of 12,591 bp, and GC content of 37.6%. Short-read transcriptome sequencing of the 91 embryo samples yielded at least 20 million transcripts per sample, apart from one sample with ~18.5 million. For haplotype 1, we annotated 43,191 genes, of which 23,572 (54.6%) had at least one functional annotation (Table 2, Figure 1). For haplotype 2, we annotated 39,675 genes, of which 22,459 (56,6%) had at least one functional annotation. Our final annotated protein set had *BUSCO* completeness scores of 93.0% for the haplotype 1 protein *fasta* file, and 94.5% for the haplotype 2 file.

### Comparison with other annelid genomes

The orthologous analysis counted 13,174 clusters in the haplotype 1 protein set. Of these clusters, 2,919 (22.2%) were unique to *Galeolaria* and 4,248 (32.2%) were shared with all three of the other annelid genomes (1738 with *Capitella* alone, 267 with *Streblospio* alone, and 253 with *Helobdella* alone; Figure 2B). The collinearity analysis detected little collinearity between genomes of *Galeolaria* and *Streblospio* (and none between their genomes and that of *Helobdella*; Figure 2C), as did our pairwise synteny search between genomes of *Galeolaria* and *Streblospio*, with only 15 matches using the default settings. Reducing the number of genes required to call synteny lead to more matches (230 alignments), but still very little synteny (Supplementary Figure S2). The *Galeolaria* genome had higher *BUSCO* completeness scores for both its assembly (95.3% *vs* 90.5% for *Streblospio*) and annotation (93.0% *vs* 61.2% for *Streblospio*), raising the possibility that unannotated genes in *Streblospio* reduced synteny detection. Assembly statistics for both genomes are reported in Supplementary Table S2.

## Discussion

To our knowledge, we have assembled the first haplotype-resolved, chromosome-level, functionally-annotated reference genome (including distinct nuclear and mitochondrial genomes) for any annelid. Haplotype-resolved assemblies like ours are emerging as key assets for genomic studies of marine invertebrates, especially broadcast spawners like *Galeolaria*, which pose unique challenges due to their high levels of heterozygosity and complex life histories (Lotterhos et al., 2021; Plough, 2016; Takeuchi et al., 2022). Accordingly, heterozygosity in the *Galeolaria* genome exceeds levels reported for other annelids (Martín-Durán et al., 2021; Zakas et al., 2022). It also exceeds levels reported for genomes of broadcast spawners in other metazoan lineages, including the pearl oyster *Pinctada fucata* (Takeuchi et al., 2022), oysters in the genus *Crassostrea* (Qi et al., 2023), and the sea star *Pisaster brevispinus* (DeBiasse et al., 2022). By capturing sequence diversity between homologous chromosomes, haplotype-resolved assemblies can offer new insights into the evolutionary origins and dynamics of such diversity across these ecologically important lineages. As such, genomic technologies that enable haplotype resolution may be essential for unlocking the full potential of marine invertebrate genomes (Takeuchi et al., 2022) and informing effective conservation and management strategies in rapidly changing marine ecosystems (Lopez et al., 2019).

Two lines of evidence (chromatin contact maps and *k*-means clustering of scaffolds) organise the nuclear genome of *Galeolaria* into 11 chromosomes, giving a diploid number of 2n=22. This is broadly consistent with cytological analyses of karyotypes in the Serpulidae (Dasgupta & Austin, 1960; Dixon et al., 1998), reporting chromosome numbers of 2n=14–28 across the family and 2n=24–26 in *Galeolaria*’s closest relatives (*Ficopomatus* and *Spirobranchus*) based on recent phylogenies (Kupriyanova et al., 2023). Variation in chromosome number among serpulids has been attributed to polyploidy followed by chromosome loss in the course of evolution (Dasgupta & Austin, 1960), and coincides with shifts in major life history traits including hermaphroditism (Kupriyanova et al., 2001). We saw no sign of sex chromosomes in *Galeolaria*, and hermaphroditism is sequential in its closest relatives (Dixon, 1981; Dixon et al., 1998) but simultaneous in more distant ones (Kupriyanova et al., 2001). New chromosome-level genomes may help resolve the genetic and evolutionary basis of this diversity, also highlighted by our comparison of *Galeolaria* with *Streblospio* (Zakas et al., 2022), *Capitella* and *Helobdella*. That these genomes show little collinearity, despite sharing a relatively large number of orthologous clusters, suggests extensive chromosomal rearrangements among them. Limited synteny between *Galeolaria* and *Streblospio*, whose genomes have similar lengths and chromosome numbers but encode distinct life histories, may likewise support such rearrangements across the sister groups they belong to (Sabellida and Spionida; Rouse et al., 2022). The only other comparison of annelid nuclear genomes reported low synteny across more distant groups (Lewin et al., 2024; Schultz et al., 2024; Zakas et al., 2022), reiterating the need for more genomes to better scrutinise these patterns.

Mitochondrial genomes are more numerous than nuclear ones, given the relative ease with which they are now assembled even from raw sequencing reads (Allio et al., 2020). The variability of annelid mitochondrial genomes, and particularly those of Serpulids, has been shown before (Struck et al., 2023; Sun et al., 2021), and is also highlighted by our alignment between the mitochondrial genomes of *Galeolaria caespitosa, Hydroides elegans*, and *Spirobranchus giganteus*. The mitochondrial genome of *Galeolaria* is similar in length to those of other serpulids, which are in turn substantially longer than reported for other annelids (Seixas et al., 2017; Sun et al., 2021). We annotated most of the 37 genes typically found in mitochondrial DNA across the Metazoa (Gissi et al., 2008; Shtolz & Mishmar, 2023), with the exception of genes for the large (*16S*) rRNA subunit and the protein *atp8*. Failure to detect these genes in *Galeolaria* could reflect their fast rates of mitochondrial sequence evolution, consistent with their variable detection in a recent comparative analysis of other serpulids (Sun et al., 2021). In particular, *atp8* is among the shortest, fastest-evolving mitochondrial genes, and has seemingly been lost in several metazoan lineages (Shtolz & Mishmar, 2023). However, it is also possible that *atp8* is erroneously reported as missing because it is highly diverged. In *Galeolaria*’s close relative *Spirobranchus giganteus*, for example, *atp8* was undetected by automated gene annotation (Seixas et al., 2017), but putatively recovered by highly sensitive manual searches at the protein level (Sun et al., 2021). As for *16S-rRNA*, it appears to be replaced in our genome by two sequences coding for *tRNA-Met* (Methionine) and one for *tRNA-Ala* (Alanine). The loss or rapid divergence of such genes in metazoans raises intriguing questions about the evolutionary drivers, and implications for mitochondrial function, that are yet to be resolved.

In summary, the high-quality genome assembly developed here for *Galeolaria* will significantly enhance the genomic resources currently available for marine invertebrates. Such resources are especially valuable for annelids, which are underrepresented in comparative genomic analyses (Shtolz & Mishmar, 2023; Zakas et al., 2022), but may deliver major insights into genome and life history evolution in an exceptionally diverse group (Martín-Zamora et al., 2023; Zakas et al., 2022) and across metazoans more broadly (Paps et al., 2023). The *Galeolaria* genome will also facilitate population genomic studies of these ecosystem engineers, whose reef-like masses of calcareous tubes enhance biodiversity on rocky shores of temperate Australia (Cole et al., 2018; Montefalcone et al., 2022). Across *Galeolaria*’s range in Australia, species have diverged and potentially recontacted in a marine biodiversity hotspot warming much faster than the global average rate (Costello, 2023; Gallegos et al., 2023; Hobday & Pecl, 2014). This system offers new scope to study climate-driven range shifts, adaptation, and diversification, and the contributions of nuclear and mitochondrial genomes to such phenomena (Gallegos et al., 2024), in a region of high endemicity vulnerable to climate change. Doing so could highlight the transformative potential of genomic tools for advancing biodiversity research and conservation efforts.

## Acknowledgements

We thank Nawar Shamaya for help with sample collection and processing, and Fisheries Victoria (RP1328) and Parks Victoria (10008784) for collection permits.

## Data availability

This Whole Genome Shotgun project has been deposited at DDBJ/ENA/GenBank under the accession JBIPTA000000000 and JBIPTB000000000.

Data generated for this study are available under NCBI BioProject PRJNA1116755. Raw sequencing data for the genome assembly (NCBI BioSample SAMN41577001) are deposited in the NCBI Short Read Archive (SRA) under SRR29259014 for PacBio HiFi sequencing data and SRR29259015 for Omni-C Illumina Short read sequencing data. Raw sequencing data for the genome annotation (NCBI BioSamples SAMN43151944 for the long reads and SAMN43151945 for the short reads) are deposited on the NCBI SRA under SRR30505187 and SRR30505186 for the long and short reads respectively. GenBank accessions for both haplotype 1 and haplotype 2 are PRJNA1116755 and PRJNA1119221. The mitochondrial genome is included in haplotype 1 (PRJNA1116755). Assembly scripts and other data for the analyses presented can be found at the following GitHub repository: https://github.com/moniquevdor/GaleolariaReferenceGenome.

