## Supplementary Material for "A phased chromosome-level genome of the annelid tubeworm *Galeolaria caespitosa*"

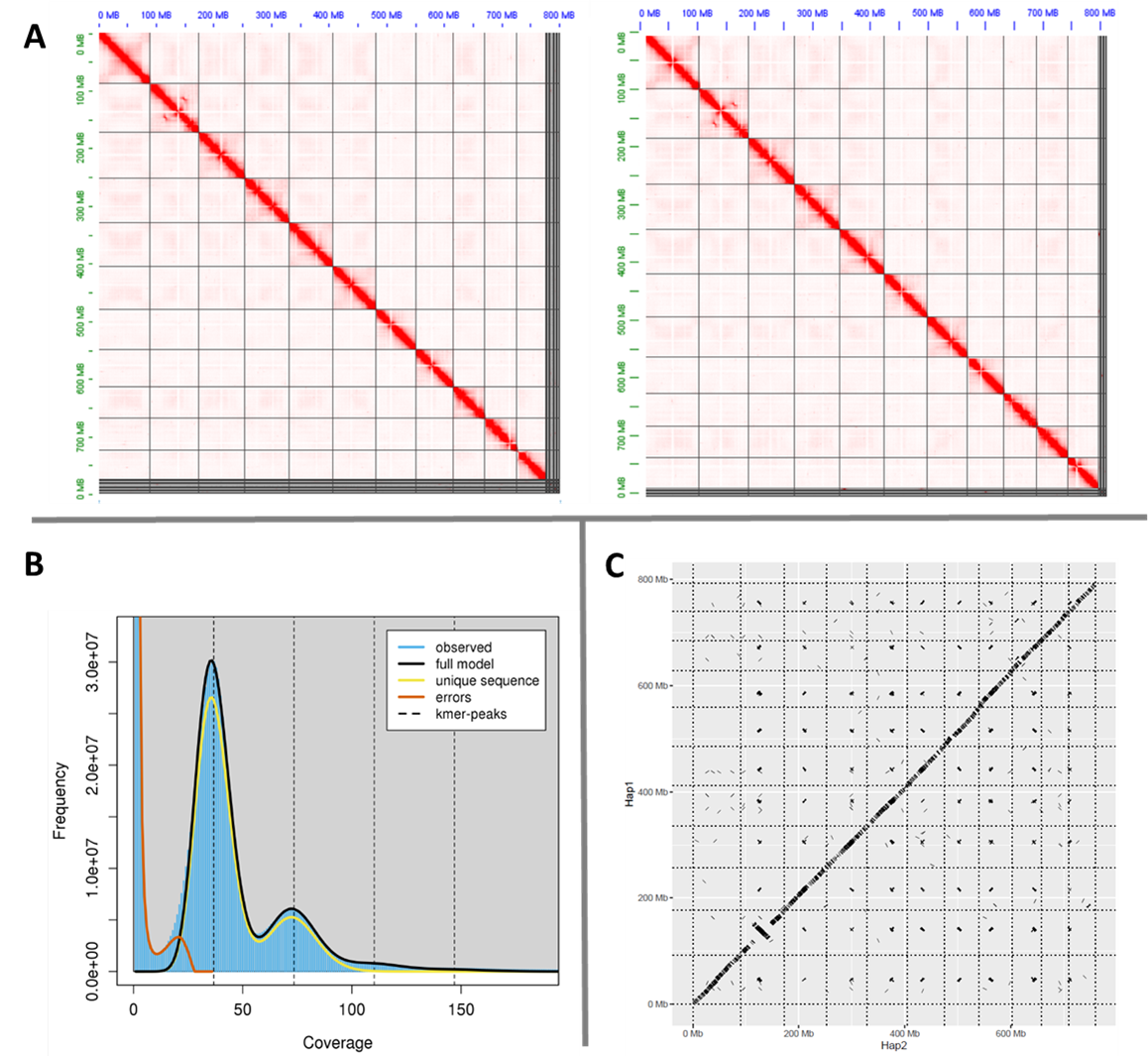

**Figure S1.** Overview of assembly metrics. **(A)** Omni-C chromatin contact maps of scaffolded genome assemblies for haplotype 1 (left) and haplotype 2 (right). Chromatin contact maps translate proximity of genomic regions in 3D space to contiguous linear organization. Each cell in the contact map corresponds to sequencing data supporting the ligation (or join) of two regions. **(B)** *K*-mer spectra generated from PacBio HiFi reads. **(C)** Dotplot alignment of haploid sets of chromosomes, with haplotype 1 on the y axis and haplotype 2 on the x axis**.**

**Table S1.** Characterization of genomic repeats from *RepeatMasker* v4.1.1 (default mode run with *rmblastn* v2.10.0+). Repeats fragmented by insertions or deletions are counted as single elements.

|  |  | number of elements | | length occupied | | percentage of sequence | |
| --- | --- | --- | --- | --- | --- | --- | --- |
|  |  | Haplotype 1 | Haplotype 2 | Haplotype 1 | Haplotype 2 | Haplotype 1 | Haplotype 2 |
| Retroelements | | 173115 | 168954 | 74221391 bp | 74468561 bp | 9.37% | 9.80% |
|  | SINEs: | 0 | 0 | 0 bp | 0 bp | 0% | 0% |
|  | Penelope | 2140 | 756 | 702048 bp | 238711 bp | 0.09% | 0.03% |
|  | LINEs: | 20264 | 16714 | 10314300 bp | 9157587 bp | 1.30% | 1.21% |
|  | CRE/SLACS | 0 | 0 | 0 bp | 0 bp | 0% | 0% |
|  | L2/CR1/Rex | 10225 | 8541 | 3903461 bp | 3754485 bp | 0.49% | 0.49% |
|  | R1/LOA/Jockey | 0 | 0 | 0 bp | 0 bp | 0% | 0% |
|  | R2/R4/NeSL | 377 | 448 | 250988 bp | 335174 bp | 0.03% | 0.04% |
|  | RTE/Bov-B | 932 | 436 | 761701 bp | 244154 bp | 0.10% | 0.03% |
|  | L1/CIN4 | 330 | 72 | 120558 bp | 66899 bp | 0.02% | 0.01% |
|  | LTR elements: | 152851 | 152240 | 63907091 bp | 65310974 bp | 8.06% | 8.60% |
|  | BEL/Pao | 16742 | 14617 | 11153105 bp | 7028094 bp | 1.41% | 0.93% |
|  | Ty1/Copia | 823 | 814 | 716056 bp | 556000 bp | 0.09% | 0.07% |
|  | Gypsy/DIRS1 | 28906 | 55303 | 17398396 bp | 27306891 bp | 2.20% | 3.60% |
|  | Retroviral | 0 | 0 | 0 bp | 0 bp | 0% | 0% |
| DNA transposons: | | 11688 | 9797 | 4741636 bp | 4391974 bp | 0.60% | 0.58% |
|  | hobo-Activator | 987 | 2460 | 505962 bp | 536505 bp | 0.06% | 0.07% |
|  | Tc1-IS630-Pogo | 190 | 0 | 65619 bp | 0 bp | 0.01% | 0% |
|  | En-Spm | 0 | 0 | 0 bp | 0 bp | 0% | 0% |
|  | MuDR-IS905 | 0 | 0 | 0 bp | 0 bp | 0% | 0% |
|  | PiggyBac | 0 | 57 | 0 bp | 16632 bp | 0% | 0% |
|  | Tourist/Harbinger | 84 | 376 | 59239 bp | 148834 bp | 0.01% | 0.02% |
|  | Other (Mirage, P-element, Transib) | 583 | 157 | 197055 bp | 42691 bp | 0.02% | 0.01% |
| Rolling-circles: | | 585 | 648 | 529853 bp | 294287 bp | 0.07% | 0.04% |
| Unclassified: | | 1031234 | 966110 | 291285589 bp | 275254969 bp | 36.76% | 36.24% |
| Total interspersed repeats: | | 1216037 | 1144861 | 370248616 bp | 354115504 bp | 46.72% | 46.62% |
| Small RNA: | | 0 | 0 | 0 bp | 0 bp | 0% | 0% |
| Satellites: | | 0 | 0 | 0 bp | 0 bp | 0% | 0% |
| Simple repeats: | | 110558 | 108128 | 7273963 bp | 7266488 bp | 0.92% | 0.96% |
| Low complexity: | | 14177 | 13538 | 668603 bp | 658256 bp | 0.08% | 0.09% |

**Table S2.** Mitochondrial genes found in (A) most metazoans, (B) *H. elegans* (the starting reference for the mitochondrial assembly), (C) *G. caespitosa* by *MITOS*, and (D) *G. caespitosa* by *MitoFinder*

| (A) Most metazoans | (B) *H. elegans* | (C) this study (*MITOS*) | (D) this study (*MitoFinder*) |
| --- | --- | --- | --- |
| COX1 | COX1 | COX1 | COX1 |
| COX2 | COX2 | COX2 |  |
| COX3 | COX3 | COX3 | COX3 |
| CYTB | CYTB | CYTB | CYTB |
| ND1 | ND1 | ND1 | ND1 |
| ND2 | ND2 | ND2 |  |
| ND3 | ND3 |  | ND3 |
| ND4 | ND4 | ND4 |  |
| ND5 | ND5 | ND5 | ND5 |
| ND6 | ND6 | ND6 |  |
| ND4L | ND4L | ND4L | ND4L |
| ATP6 | ATP6 | ATP6 |  |
| ATP8 |  |  |  |
| 12S rRNA | 12S rRNA | 12S rRNA |  |
| 16S rRNA | 16S rRNA |  |  |
| tRNA-Ala | tRNA-Ala | tRNA-Ala | tRNA-Ala |
| tRNA-Arg | tRNA-Arg | tRNA-Arg | tRNA-Arg |
| tRNA-Asn | tRNA-Asn | tRNA-Asn | tRNA-Asn |
| tRNA-Asp | tRNA-Asp | tRNA-Asp | tRNA-Asp |
| tRNA-Cys | tRNA-Cys | tRNA-Cys | tRNA-Cys |
| tRNA-Gln | tRNA-Gln | tRNA-Gln | tRNA-Gln |
| tRNA-Glu | tRNA-Glu | tRNA-Glu | tRNA-Glu x4 |
| tRNA-Gly | tRNA-Gly | tRNA-Gly | tRNA-Gly x2 |
| tRNA-His | tRNA-His | tRNA-His | tRNA-His |
| tRNA-Ile | tRNA-Ile | tRNA-Ile | tRNA-Ile |
| tRNA-Lys | tRNA-Lys | tRNA-Lys | tRNA-Lys |
| tRNA-Met | tRNA-Met | tRNA-Met x5 | tRNA-Met x5 |
| tRNA-Phe | tRNA-Phe | tRNA-Phe | tRNA-Phe |
| tRNA-Pro | tRNA-Pro | tRNA-Pro | tRNA-Pro |
| tRNA-Thr | tRNA-Thr | tRNA-Thr | tRNA-Thr |
| tRNA-Trp | tRNA-Trp | tRNA-Trp | tRNA-Trp |
| tRNA-Tyr | tRNA-Tyr | tRNA-Tyr | tRNA-Tyr |
| tRNA-Val | tRNA-Val | tRNA-Val | tRNA-Val |
| tRNA-Leu x2 | tRNA-Leu x2 | tRNA-Leu x2 | tRNA-Leu x2 |
| tRNA-Ser x2 | tRNA-Ser x2 | tRNA-Ser x2 | tRNA-Ser x2 |

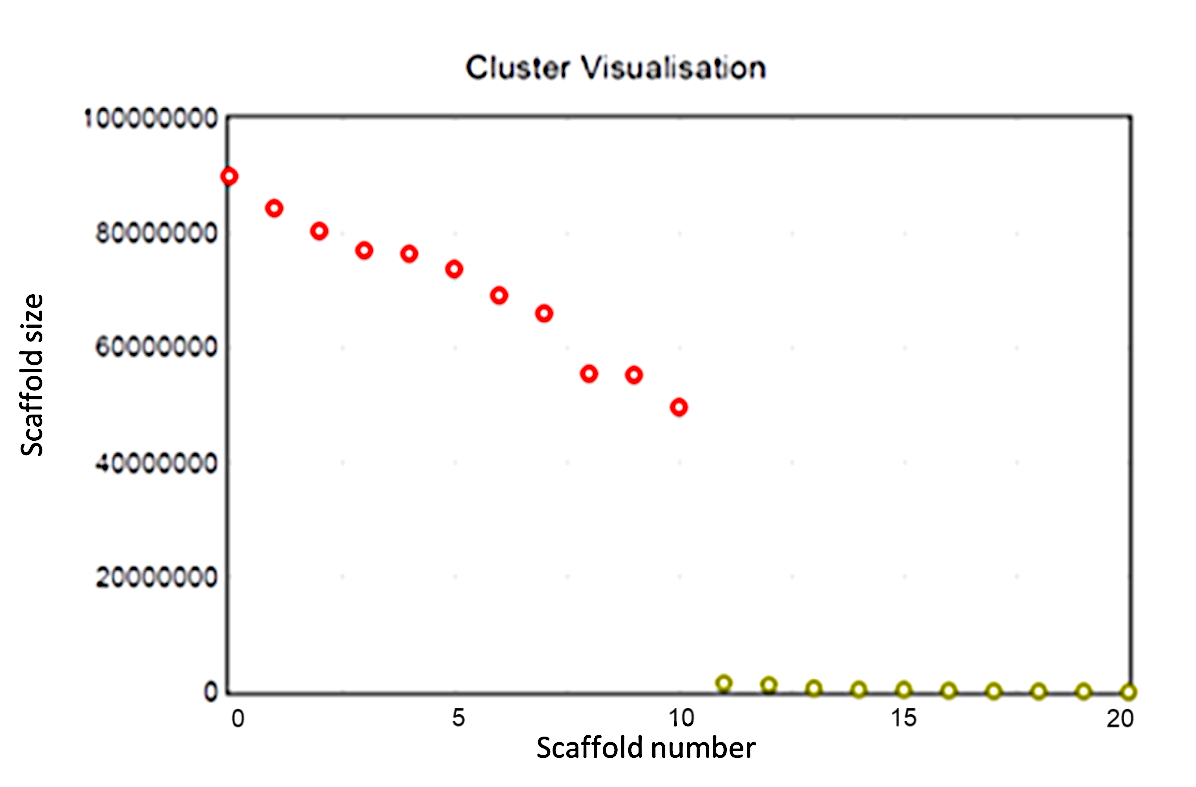

**Figure S2.** *K*-means clustering performed on the twenty longest scaffolds in the *Galeolaria caespitosa* genome. Red circles indicate scaffolds clustered in group 1 and green circles indicate scaffolds clustered in group 2.

**Table S3.** BUSCO metrics for *Galeolaria caespitosa* and *Streblospio benedicti* against the eukaryote ortholog database.

|  | *Galeolaria* *caespitosa* primary assembly (scaffold-level) | *Streblospio benedicti* (Zakas et al., 2022) |
| --- | --- | --- |
| Number of scaffolds | 583 | 6,112 |
| Total length (Mbp) | 803.5 | 701.4 |
| Largest scaffold (Mbp) | 90.0 | 64.9 |
| Scaffold N50 (bp) | 76.5 | 56.1 |
| Scaffold L50 | 5 | 6 |
| GC (%) | 34.5% | 37.9% |
| Assembly BUSCO scores |  |  |
| Complete BUSCOs (C) | 95.3% | 90.5% |
| Complete and single-copy BUSCOs (S) | 94.3% | 87.8% |
| Complete and duplicated BUSCOs (D) | 0.4% | 2.7% |
| Fragmented BUSCOs (F) | 4.3% | 6.3% |
| Missing BUSCOs (M) | 0.4% | 3.2% |
| Annotation BUSCO scores |  |  |
| Complete BUSCOs (C) | 93.0% | 61.2% |
| Complete and single-copy BUSCOs (S) | 90.6% | 56.5% |
| Complete and duplicated BUSCOs (D) | 2.4% | 4.7% |
| Fragmented BUSCOs (F) | 5.9% | 9.4% |
| Missing BUSCOs (M) | 1.1% | 29.4% |

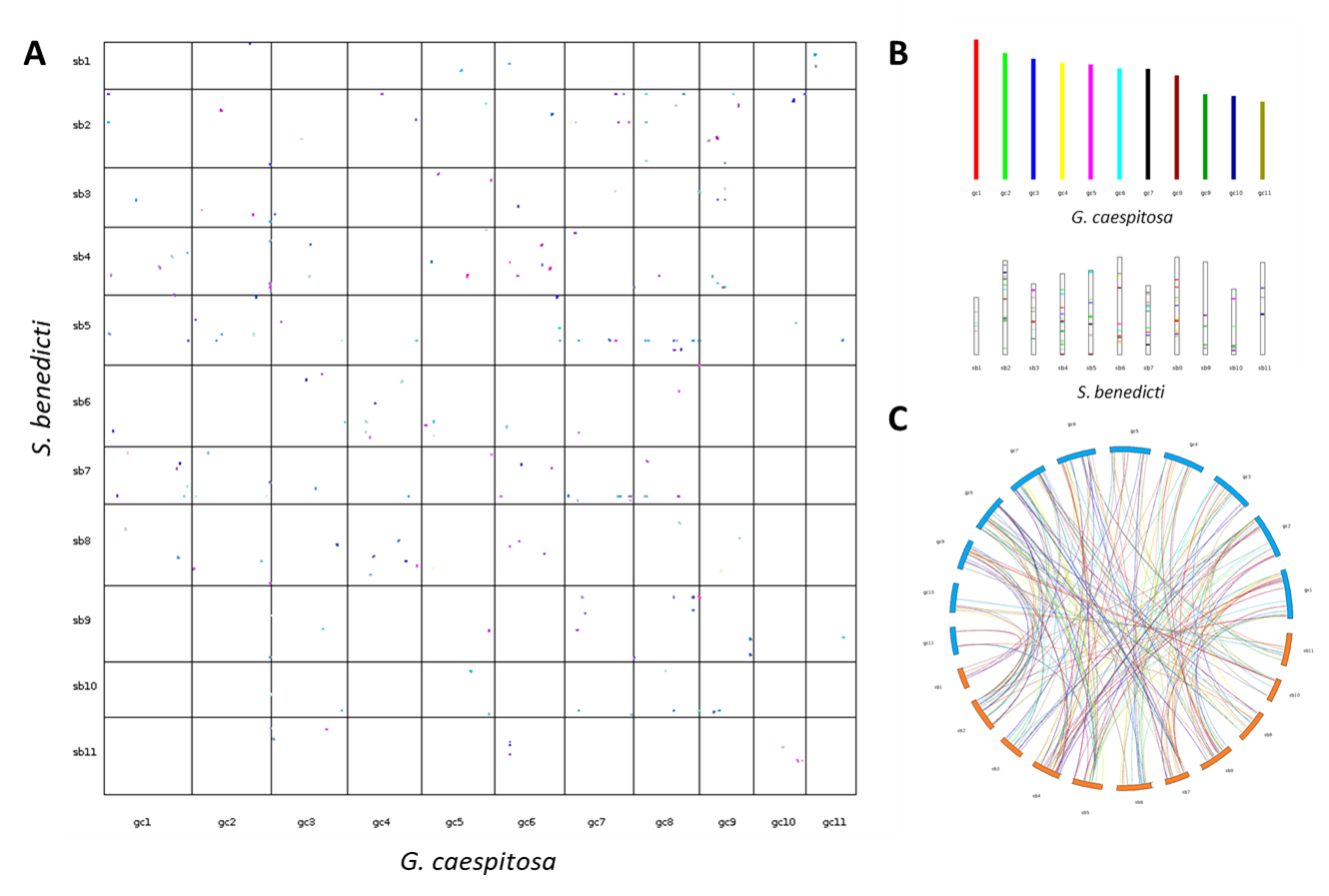

**Figure S3.** Results of pairwise synteny search from *MCScanX*. (A) Dot plot showing regions of synteny between genomes of *G. caespitosa* and *S. benedicti*. Axes denote genes on the 11 chromosomes of each species, and coloured dots denote genomic regions of at least 3 genes shared between species. (B) Bar plot showing estimated proportions of sequences present in *G. caespitosa* (top) and *S. benedicti* (bottom). The corresponding sequences from *G. caespitosa* in *S.* benedicti are indicated through matching colours. (C) Circle plot showing genomic regions that are collinear between *G. caespitosa* (blue) and *S. benedicti* (orange).
